# Enhancing acute kidney injury regeneration by promoting cellular dedifferentiation in zebrafish

**DOI:** 10.1101/434951

**Authors:** Lauren Brilli Skvarca, Hwa In Han, Eugenel B. Espiritu, Maria A. Missinato, Elizabeth R. Rochon, Michael D. McDaniels, Beth L. Roman, Joshua S. Waxman, Simon C. Watkins, Alan J. Davidson, Michael Tsang, Neil A. Hukriede

**Affiliations:** Department of Developmental Biology, University of Pittsburgh, Pittsburgh, PA, USA; Department of Pathology, University of Pittsburgh, Pittsburgh, PA, USA; Pittsburgh Heart, Lung, and Blood Vascular Medicine Institute, Department of Medicine, University of Pittsburgh, Pittsburgh, PA, USA; Human Genetics, Graduate School of Public Health, University of Pittsburgh, Pittsburgh, PA, USA; Heart Institute, Molecular Cardiovascular Biology Division, Cincinnati Children’s Hospital Medical Center, Cincinnati OH, USA; Department of Cell Biology and Center for Biological Imaging, University of Pittsburgh, Pittsburgh, PA, USA; Center for Critical Care Nephrology, University of Pittsburgh, Pittsburgh, PA, USA; Molecular Medicine and Pathology, University of Auckland, Auckland, New Zealand; Sanford Burnham Prebys Medical Discovery Institute, La Jolla, California USA

**Keywords:** acute kidney injury, macrophages, renal proximal tubule cell, dedifferentiatio, HDAC inhibitor, therapeutic

## Abstract

Acute kidney injury (AKI) is a serious disorder for which there is no approved pharmaceutical treatment. Following injury, native nephrons display limited regenerative capabilities, relying on the dedifferentiation and proliferation of renal tubular epithelial cells (RTECs) that survive the insult. Previously, we identified 4-(phenylthio)butanoic acid (PTBA), a histone deacetylase inhibitor (HDI) that enhances renal recovery and showed that PTBA treatment increased RTEC proliferation and reduced renal fibrosis. Here, we investigated the regenerative mechanisms of PTBA in zebrafish models of larval renal injury and adult cardiac injury. With respect to renal injury, we showed that delivery of PTBA using an esterified prodrug (UPHD25) increases the reactivation of the renal progenitor gene Pax2a, enhances dedifferentiation of RTECs, reduces Kidney injury molecule-1 expression, and lowers the number of infiltrating macrophages. Further, we find that the effects of PTBA on RTEC proliferation depend upon retinoic acid signaling and demonstrate the therapeutic properties of PTBA are not restricted to the kidney but also increase cardiomyocyte proliferation and decrease fibrosis following cardiac injury in adult zebrafish. These studies provide key mechanistic insights into how PTBA enhances tissue repair in models of acute injury and lay the groundwork for translating this novel HDI into the clinic.

**SUMMARY STATEMENT:** Mortality associated with acute kidney injury (AKI) is in part due to limited treatments available to ameliorate kidney injury. We identified a compound that enhances AKI recovery by promoting cellular dedifferentiation.

## INTRODUCTION

Acute kidney injury (AKI) is a rapid decline in renal function that results in 2 million deaths annually worldwide and accounts for billions in US healthcare costs (Chertow et al., 2005; Murugan and Kellum, 2011). AKI frequently occurs in a hospitalized setting where 3-20% of patients are affected (Fang et al., 2010; Uchino et al., 2006). It is even more common in the intensive care unit, with up to 67% of patients affected (Bagshaw et al., 2008; Cruz et al., 2007; Hoste et al., 2006; Ostermann and Chang, 2007). It is now recognized that AKI may not be an isolated event; rather, many patients who recover clinically do not regain baseline renal function and are at increased risk for developing chronic kidney disease (Chawla and Kimmel, 2012; Coca et al., 2012; Kjellstrand et al., 1981; Liano et al., 2007; Venkatachalam et al., 2010).

AKI represents a disease spectrum with numerous contributing causes (Levey and James, 2017). Since renal biopsies are not often performed in AKI patients, the underlying physiology and histopathology have been largely defined using rodent models (Lieberthal and Nigam, 2000; Murugan and Kellum, 2011). These studies have identified key cellular players during AKI events, including roles for the tubular epithelium and the immune system. Specifically, damaged renal tubular epithelial cells (RTECs) undergo dedifferentiation to a mesenchymal state by reactivating pathways common during early renal development (Benigni et al., 2010; Cirio et al., 2014; Imgrund et al., 1999; Lin et al., 2010; Terada et al., 2003; Villanueva et al., 2006; Witzgall et al., 1994). These surviving RTECs proliferate and repopulate areas of lost cells in the nephron (Humphreys et al., 2008). Additionally, injured RTECs activate an immune response by increased expression of toll-like receptors and production of pro-inflammatory cytokines to attract leukocytes to the kidney (Wolfs et al., 2002; Wu et al., 2007). These signals result in the rapid activation of the innate immune system, with neutrophils arriving within 30 minutes of ischemia reperfusion (IR-AKI) injury (Awad et al., 2009; Kelly et al., 1996; Li et al., 2008; Rabb et al., 1994; Thornton et al., 1989). In addition, heterogeneous macrophage populations participate in both tissue injury and repair (Cao et al., 2015; Ricardo et al., 2008). Early in the course of renal injury, pro-inflammatory macrophages (called either classically activated or M1) infiltrate and perpetuate damage by secreting pro-inflammatory cytokines and producing reactive oxygen species (ROS) (Cao et al., 2015; Jo et al., 2006; Lee et al., 2011). Subsequently, phagocytosis of cellular debris and other environmental cues trigger suppression of M1 macrophages and promote polarization toward a reparative macrophage phenotype (called either alternatively activated or M2) (Cao et al., 2015; Lee et al., 2011). M2 macrophages are generally considered to promote tissue repair, proliferate *in situ* (Zhang et al., 2012), and protect RTECs from apoptosis, and promote cell cycle progression (Lin et al., 2010; Schmidt et al., 2013; Sola et al., 2011). These coordinated responses are critical for nephron repair and recovery from AKI (Han et al., 2018).

Despite advancements in understanding AKI pathophysiology, there is an unmet need for clinical therapies. No targeted clinical treatments are currently available that accelerate renal recovery or decrease fibrosis when administered after injury. We previously identified 4-(phenylthio)butanoic acid (PTBA), a novel short-chain carboxylic acid class histone deacetylase inhibitor (HDI), which are considered HDAC class I specific (de Groh et al., 2010; Fass et al., 2011). We previously evaluated the therapeutic potential of PTBA, delivered as a prodrug of either a methyl ester (UPHD25) or amide (UPHD186) (Cianciolo Cosentino et al., 2013; Skrypnyk et al., 2016). In zebrafish and mouse models of AKI, we have shown that PTBA enhances survival, increases RTEC proliferation, ameliorates injury, and reduces renal scarring (Cianciolo Cosentino et al., 2013; Novitskaya et al., 2014; Skrypnyk et al., 2016). Importantly, PTBA was efficacious when delivered post-AKI in all animal models, heightening the potential for clinical applicability.

In this study, we utilized the optical transparency of zebrafish larvae to characterize the cellular mechanisms by which PTBA enhances the regenerative response following AKI. Since we have previously shown that PTBA exhibits similar efficacy in both zebrafish and mammalian AKI models, here we focus on zebrafish AKI studies, as we can easily and precisely deliver the compound to zebrafish larvae and subsequently track cell populations by both static and live imaging methods. We demonstrate that delivering PTBA as a prodrug (UPHD25) increases RTEC dedifferentiation, attenuates tubular injury, lowers total macrophage recruitment, and decreases the number of inflammatory (M1) macrophages. We found that the pro-regenerative effects of PTBA depend on intact retinoic acid (RA) signaling, in line with HDACs being modulators of RA signaling and this pathway playing a key early role in kidney regeneration (Brilli et al., 2013; Chiba et al., 2015). Finally, we looked at the applicability of PTBA to enhance cardiac repair, another RA-dependent model of regeneration (Kikuchi et al., 2011), and found that treatment promotes cardiomyocyte proliferation and decreases fibrosis. Overall, these studies provide insight into the cellular mechanism underlying PTBA-driven regeneration.

## RESULTS

### PTBA increases RTEC dedifferentiation and repair responses post-AKI

We have previously shown that PTBA treatment increases RTEC proliferation in a zebrafish larval model of AKI (Cianciolo Cosentino et al., 2013; Hentschel et al., 2005). To assess whether PTBA increases the population of cells that drive AKI mediated proliferation, we examined whether UPHD25 treatment increases Pax2a expression after renal injury, since Pax2a (an early acting renal transcription factor) is a well-known marker of gene reactivation during RTEC dedifferentiation (Humphreys et al., 2011; Imgrund et al., 1999; Maeshima et al., 2002; Villanueva et al., 2006). Larvae were injected with gentamicin (gent-AKI) between 78-82 hours post-fertilization (hpf), followed by single-dose UPHD25 treatment at 2 days post injection (dpi). To identify proximal tubules (PT), we used the *Tg(PT:EGFP)* line (Cianciolo Cosentino et al., 2013). In gent-AKI larvae, Pax2a is reactivated in RTECs within 48 hrs after gentamicin injection and expression is maintained in a population of RTECs through at least 3 dpi (Figure 1A-C). When gent-AKI larvae are treated with UPHD25, the number of Pax2a-positive RTECs significantly increases (Figure 1D,E). To demonstrate that Pax2a is expressed in proliferating cells in zebrafish larvae in AKI, as has been shown in murine models (Humphreys et al., 2011), we co-stained for Pax2a and proliferating cell nuclear antigen (PCNA), which marks proliferating cells in S phase. PCNA co-localized with Pax2a in both UPHD25 and DMSO-treated larvae (Figure 1F,G).

**Figure 1.**
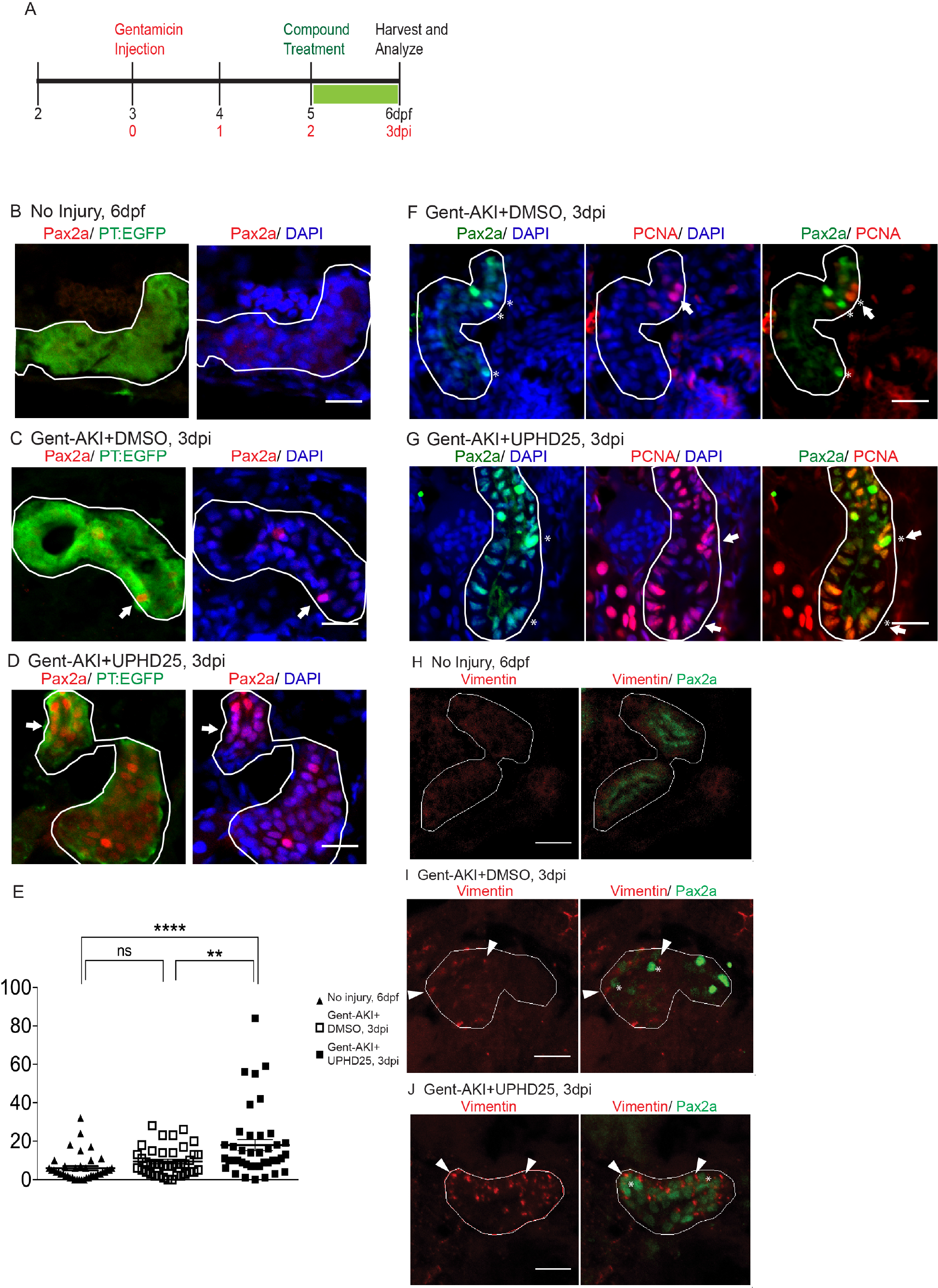
UPHD25 treatment increases Pax2a reactivation and aids in dedifferentiation during AKI. (A) Experiment schematic: *Tg(PT:EGFP)* larvae were injected with gentamicin at 3 dpf to induce AKI (gent-AKI). At 2 dpi, gent-AKI larvae were treated with 1% DMSO or 1 µM UPHD25 for 24 hrs (2-3 dpi) then harvested for analysis. (B-D) Immunofluorescence staining of Pax2a (red), proximal tubule (PT, green), and nuclei (DAPI, blue) of age-matched 6 dpf uninjured (B), 3 dpi gent-AKI+DMSO (C), and 3 dpi gent-AKI+UPHD25 (D) larvae. Nuclear localization of Pax2a was shown by overlaying with nuclear counterstain, DAPI (blue). PT is outlined in white and Pax2a^+^ RTECs are marked with arrows. (E) Quantification of Pax2a^+^ cells. Mean_gent-_ AKI+DMSO=9.5, N=40, vs. mean_gent-AKI+UPHD25_=18.0, N=40. Data pooled from 3 biological replicates are shown expressed as mean+/-SEM. One-way ANOVA: **p<0.01, ****p<0.001, n.s. = not significant. (F-G) Immunofluorescence co-stain of Pax2a (green), PCNA (red), and nuclei (DAPI, blue) in gent-AKI+DMSO (F) and gent-AKI+UPHD25 (G). Nuclear localization of PCNA was shown by overlaying with nuclear counterstain, DAPI (blue). Pax2a+ are marked with asterisks and PCNA+ are marked with arrows. (H-J) Immunofluorescence co-stain of Vimentin (red, cytosolic) and Pax2a (green) in PT of age-matched 6 dpf uninjured (H), 3 dpi gent-AKI+DMSO (I), and 3 dpi gent-AKI+UPHD25 (J). Pax2a+ are marked with asterisks and Vimentin+ RTECs are marked with arrowheads. PT is outlined in white. Scale bar= 20 µm.

To confirm that Pax2a reactivation is associated with dedifferentiation of RTECs, we examined the effect of PTBA on Vimentin expression, an intermediate filament protein that increases in dedifferentiated RTECs after AKI (Witzgall et al., 1994). We treated gent-AKI *Tg(PT:EGFP)* fish with DMSO or UPHD25 at 2 dpi and examined co-expression of Vimentin and Pax2a. While RTECs in uninjured control larvae did not show Vimentin nor Pax2a staining (Figure 1H), RTECs in gent-AKI larvae treated with either DMSO or UPHD25 showed positive Vimentin expression that is associated with Pax2a-positive cells (Figure 1I,J).

Since treatment increases the dedifferentiation and proliferation (Cianciolo Cosentino et al., 2013) of proximal tubule cells, we examined the effect of PTBA on levels of kidney injury molecule-1 (Kim-1) expression, which increases in RTECs after AKI and also plays a role in fibrosis and leukocyte recruitment (Humphreys et al., 2013; Ichimura et al., 1998). Following gent-AKI in zebrafish larvae, robust Kim-1 staining is detectable in injured fish by 2 dpi (Chiba et al., 2015). To quantify Kim-1 protein, we measured the percent area of fluorescence within the tubule (Figure 2A-C). Compared to gent-AKI+DMSO, UPHD25-treated fish showed a decrease in renal Kim-1 signal at 3 dpi (Figure 2D). Collectively, these data demonstrate that Pax2a reactivation occurs in dedifferentiated RTECs after gentamicin injury, and that PTBA mitigates cellular injury in RTECs.

**Figure 2.**
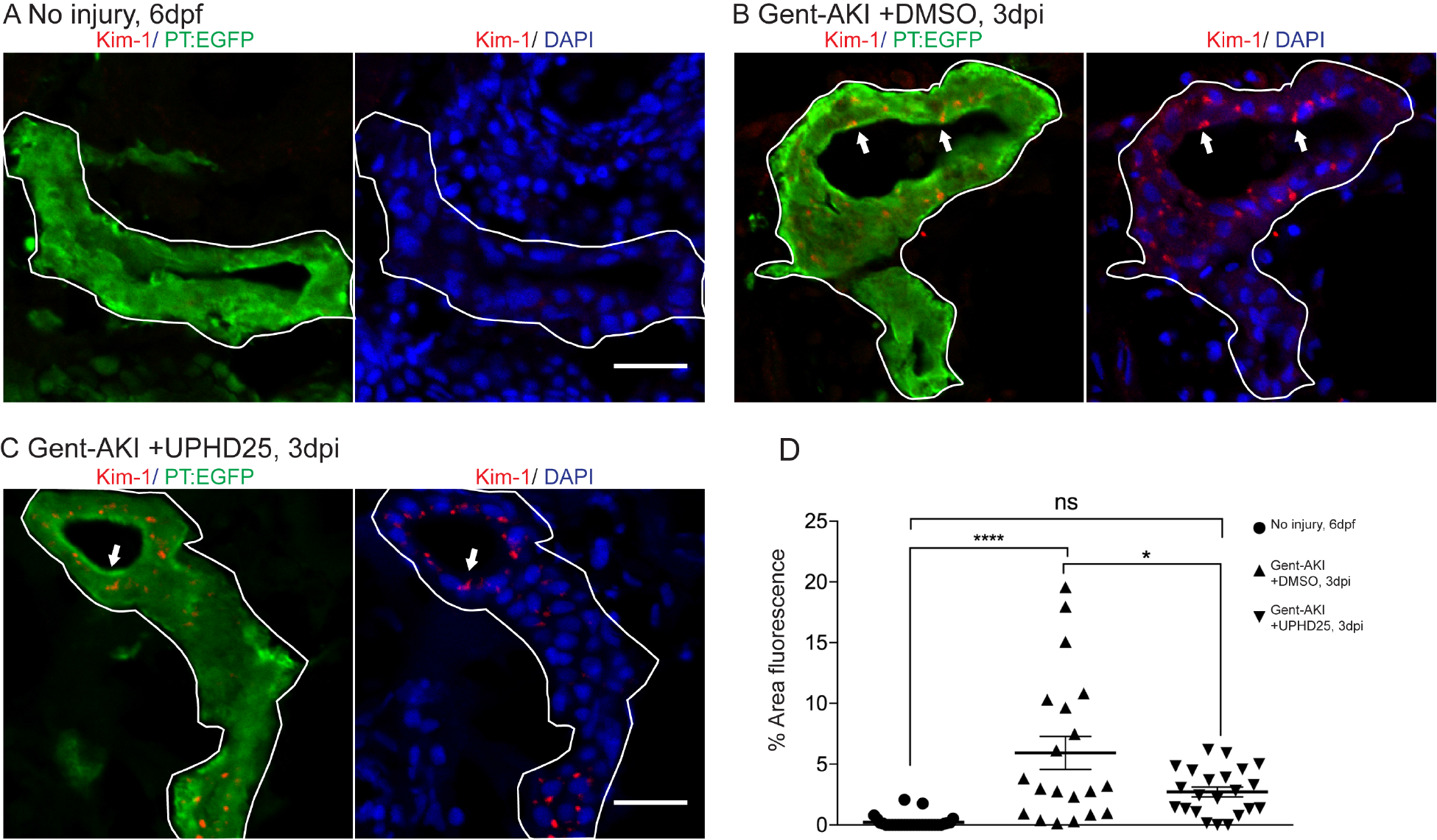
UPHD25 treatment decreases Kim-1 expression level after AKI. (A) Immunofluorescence of Kim-1 expression in 6dpf uninjured (A) 3 dpi gent-AKI +DMSO (B), and gent-AKI +UPHD25 (C) larvae. Apical localization of Kim-1 was shown by overlaying with nuclear counterstain, DAPI (blue). Histological sectioning poses a challenge to obtaining perfectly perpendicular transversal cut to observing Kim-1 at apical localization. See (C) to see an ideal perpendicular transversal section to observe apical expression of Kim-1. PT are outlined in white and RTECs with Kim-1 expression are marked with arrows. (D) Quantification of Kim-1 was acquired via area of Kim-1 expression in PT. Mean_No injury_= 0.22% (N=29) vs. mean_gent-AKI_+DMSO=5.92% (N=20) vs. mean_gent-AKI +UPHD25_=2.71% (N=22). Data pooled from 3 biological replicates are shown expressed as mean+/-SEM. One-way ANOVA: *p<0.05, ****p<0.001, n.s.=not significant. Scale bar= 20 µm.

### Immune system response in zebrafish gent-AKI

Activation of Kim-1 in RTECs has been associated with prolonged inflammatory response and fibrosis in mice (Humphreys et al., 2013). Thus, we determined whether PTBA affects the leukocyte response in the zebrafish gentamicin-AKI model. In mammals, AKI results in the rapid influx of neutrophils and macrophages (Cao et al., 2015; Jo et al., 2006; Lee et al., 2011; Li et al., 2008; Ricardo et al., 2008), but these responses have not yet been characterized during AKI in zebrafish. Since the innate immune response is functional in zebrafish larvae by 3 dpf (Keightley et al., 2014), we first determined whether neutrophils and macrophages also infiltrate the pronephric kidney after gent-AKI. To evaluate the neutrophil response, we performed time-lapse confocal imaging in gent-AKI transgenic fish expressing mCherry driven by the *cadherin*-17 promoter, *Tg(cdh17:mCherry),* a renal tubule marker (Sanker et al., 2013), and enhanced GFP driven by the *lysozyme C* promoter, *Tg(lyz:EGFP),* a marker of neutrophils (Figure 3A) (Ellett et al., 2011; Kitaguchi et al., 2009). We captured z-stack images of the PT region over 17 hrs beginning at 24 hpi. In uninjured fish, EGFP+ neutrophils move rapidly with few cells accumulating near the PT (Figure 3B and Suppl. Movie 1). In contrast, neutrophils in gent-AKI fish move slowly, and several migrated adjacent to the mCherry-positive kidney epithelium (Figure 3B and Suppl. Movie 2). In order to quantify the response, we acquired samples at several time points and quantified EGFP+ neutrophils adjacent to the PT by examining serial sections (Figure 3C). At 4 dpf (1 dpi), there is no change in the number of renal neutrophils after gentamicin injection (Figure 3D). However, gent-AKI larvae showed significantly more renal neutrophils at both 5 dpf (2 dpi) and 6 dpf (3 dpi), compared to uninjured controls (Figure 3D).

**Figure 3.**
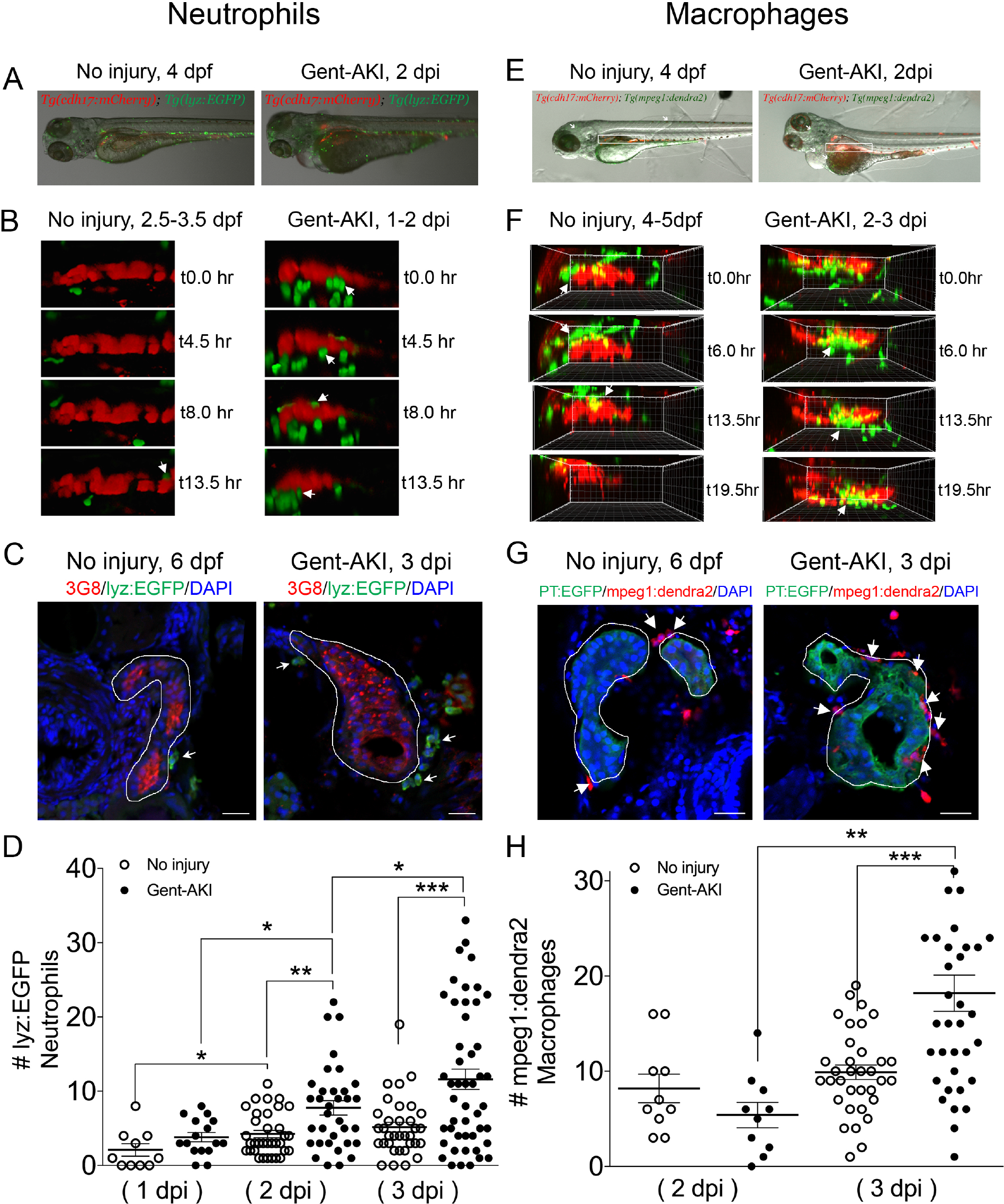
Neutrophil and macrophage populations change in the kidney field after AKI. (A-D) *Tg(chd17:mCherry); Tg(lyz:EGFP)* transgenic zebrafish were used for neutrophil analyses. (E-H) *Tg(chd17:mCherry); Tg(mpeg1;dendra2)* transgenic zebrafish were used for macrophage analyses. (A, E) Transgenic lines were injected with gentamicin at 3 dpf and imaged at 2 dpi (B, F) Snapshots of live imaging of *lyz*+ neutrophils imaged at 1 dpi for 13.5 hrs (B) and *mpeg1*+ macrophages imaged at 2 dpi for 19.5 hrs (F) in no injury and gent-AKI setting. (C, G) Immunofluorescence co-stain of PT (red or green) and neutrophils (green) (C) or macrophages (red) (G) in no-injury and gent-AKI at 3 dpi. PT is outlined in white and adjacent leukocytes are marked with arrows. (D, H) Quantification of neutrophil (D) and macrophage (H) numbers adjacent to PT before and after injury. Mean_No injury 5dpf_ =4.24 (N=34) vs. mean_2dpi_ =7.76 (N=34) vs. mean_No injury 6dpf_ =5.15 (N=34) vs. mean_3dpi_=11.60 (N=48). Adjacent leukocytes were counted for both no injury and gent-AKI at 1, 2, 3 dpi (D) and 2, 3 dpi (H). Data pooled from 3 biological replicates are shown expressed as mean+/-SEM. One-way ANOVA: *p<0.05, **p<0.01, ***p<0.005. Scale bar=20 µm.

We performed similar imaging and histology studies to evaluate the macrophage response. To visually detect macrophages, we utilized the transgenic reporter line, *Tg(mpeg1:dendra2)* in which macrophages are green (Harvie et al., 2013). We performed live time-lapse imaging in *Tg(cdh17:mCherry); Tg(mpeg1:dendra2)* double transgenic fish after gentamicin injection (Figure 3E). We captured z-stack images of the PT region over 20 hrs beginning at 48 hpi and observed an influx of macrophages to the PT in gent-AKI fish. Uninjured fish display dendra2+ macrophages circulating rapidly; however, only a few macrophages make prolonged contact with the PT (Figure 3F and Suppl. Movie 3). In contrast, gent-AKI fish show recruitment and retention of many macrophages to the mCherry-positive kidney epithelium (Figure 3F and Suppl. Movie 4). To quantify this response, we fixed *Tg(mpeg1:dendra2); Tg(PT:EGFP)* double transgenic fish and counted the number of dendra2-positive macrophages adjacent to the PT (Figure 3G). Compared to uninjured fish, gent-AKI larvae showed an increased number of renal macrophages at 3 dpi (Figure 3G,H). Taken together, these data indicate that the gent-AKI zebrafish larvae show a robust innate immune response between 2-3 dpi.

### Effect of PTBA on the immune system

Having characterized the timing of neutrophil and macrophage influx during gent-AKI in zebrafish larvae, we assessed whether PTBA affects the innate immune response. We treated gent-AKI *Tg(lyz:EGFP)* or *Tg(mpeg1:dendra2)* with DMSO or UPHD25, and quantified renal neutrophils and macrophages at 3 dpi (Figure 4A,B). At this time point, there was no significant change in the number of neutrophils, but there was a small but significant decrease in the number of macrophages between treatment groups. This suggests that UPHD25 treatment does not affect initial neutrophil recruitment but may decrease the overall number of macrophages that are recruited to the PT (Figure 4C,D).

**Figure 4.**
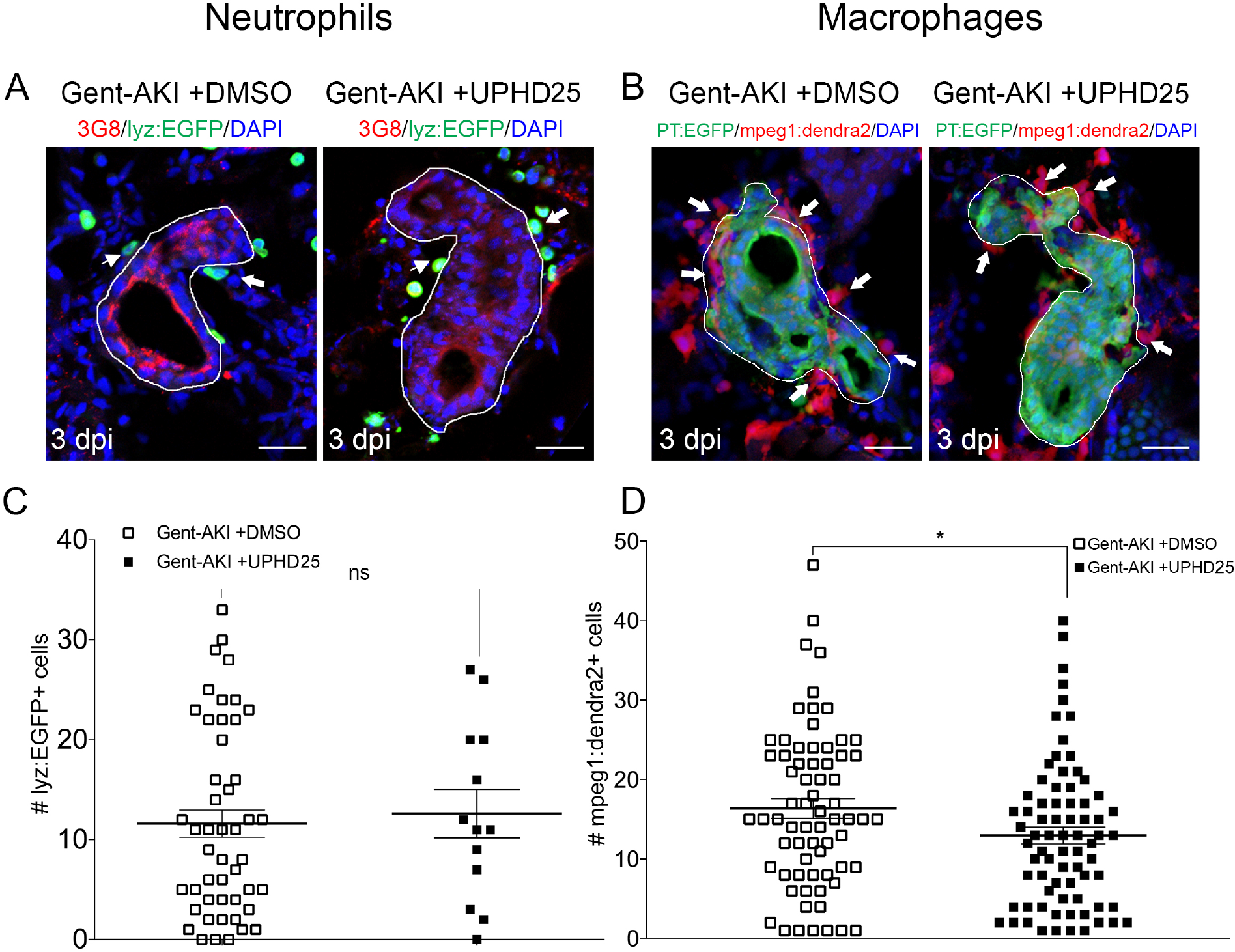
UPHD25 treatment has no effect on neutrophil response but lowers total macrophage recruitment during early AKI phase. (A, C) *Tg(chd17:mCherry); Tg(lyz:EGFP)* transgenic zebrafish were used for neutrophil analyses. (B, D) *Tg(chd17:mCherry); Tg(mpeg1;dendra2)* transgenic zebrafish were used for macrophage analyses. (A, B) Immunofluorescence co-stain of PT (red or green) and neutrophils (green) (A) and macrophages (red) (B) in gent-AKI+DMSO and gent-AKI +UPHD25. PT is outlined in white and leukocytes are marked with arrows. (C) Quantification of neutrophil numbers showed no significance between DMSO and UPHD25 treatment groups. (D) Quantification of macrophage numbers showed significant decrease in macrophage numbers in UPHD25 treatment group. Mean_gent-AKI_+DMSO=16.38 N=69 vs. mean_gent-AKI +UPHD25_=12.99 N=75. Data pooled from 3 biological replicates are shown expressed as mean+/-SEM. 2 tailed t-test: *p<0.05, n.s.= not significant. Scale bar=20µm.

Since the mpeg1 transgenic line marks multiple macrophage phenotypes, we wanted to determine whether UPHD25 treatment affected macrophage polarization (Cao et al., 2015; Jo et al., 2006; Lee et al., 2011). Studies have shown that zebrafish undergo M1/M2 polarization akin to their mammalian counterparts (Nguyen-Chi et al., 2015; Wiegertjes et al., 2016). Therefore, we utilized the *Tg(mpeg1:dendra2)* line and stained for TNFα, an M1-specific pro-inflammatory cytokine (Figure 5A-C) (Nelson et al., 2013). Since other pro-inflammatory cells also express TNFα, we quantified the ratio of cells that co-labelled TNFα+/mpeg1:dendra2+. In gent-AKI+DMSO treated fish, many M1 macrophages are recruited to RTECs, while gent-AKI+UPHD25 treated fish display fewer M1 macrophages (Figure 5A-C). To quantify M2, macrophages, we utilized the same method, but used an M2 specific enzyme, Arginase-2 (Suppl. Figure 1) (Wiegertjes et al., 2016). Comparing gent-AKI+DMSO with gent-AKI+UPHD25 treated fish, we found no significant change in the number of M2 macrophages in the renal field (Figure 5D-F). Overall, these results suggest that PTBA reduces total macrophage recruitment and the number of inflammatory macrophages to the damaged tubule.

**Figure 5.**
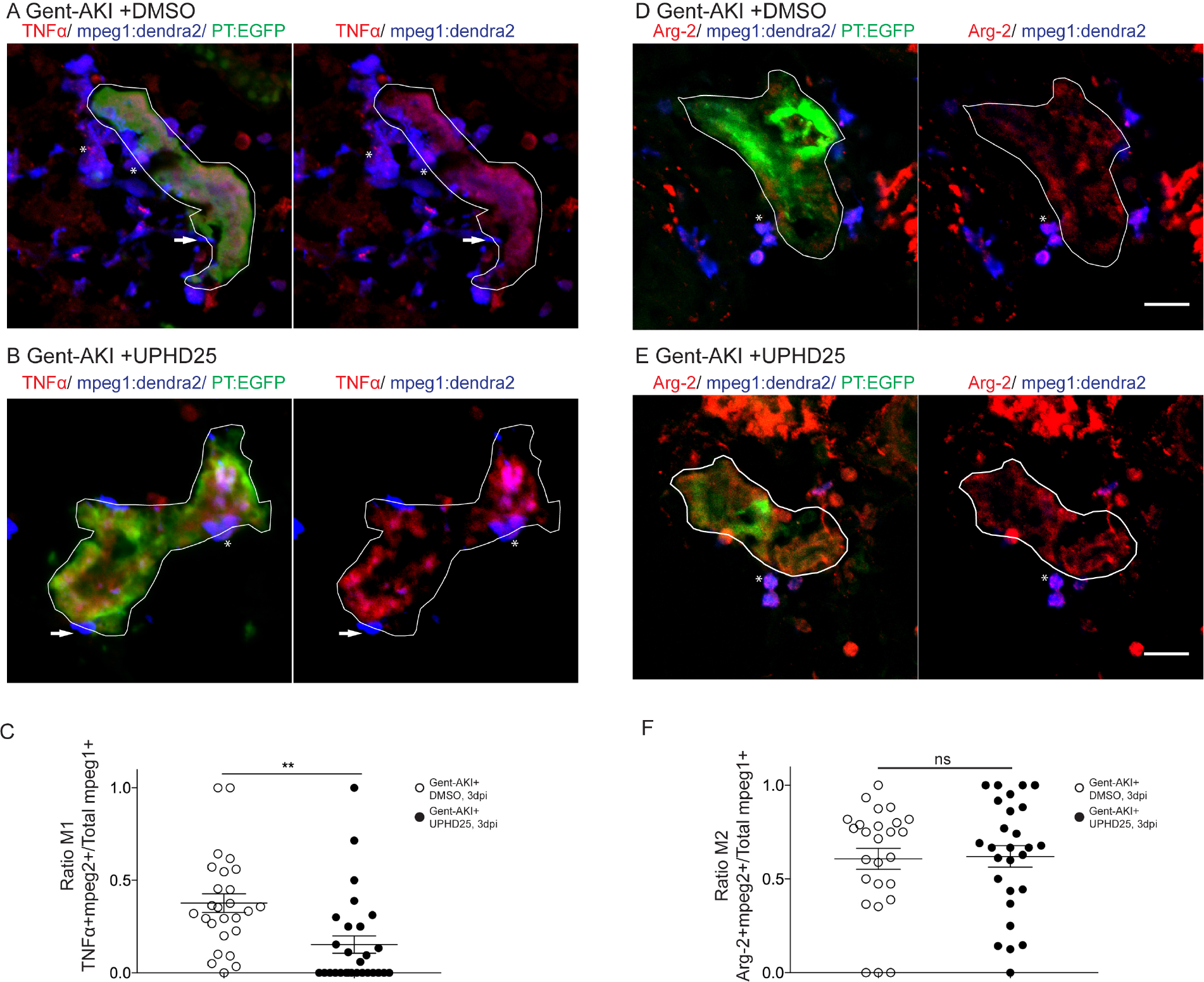
UPHD25 treatment reduces total M1 macrophage population, but not M2 macrophage population. *Tg(mpeg1:dendra2); Tg(PT:EGFP)* transgenic zebrafish were used for macrophage polarization analysis. (A, B) Immunofluorescence co-stain of TNFα (red), macrophages (blue), and PT (green) in gent-AKI +DMSO (A) and gent-AKI +UPHD25 (B). PT is outlined in white. TNFα+/mpeg1+ are marked with an asterisk and TNFα-/mpeg1 are marked with an arrow. (C) Quantification of M1 macrophage recruitment by counting TNFα+/mpeg1+ cells adjacent to PT. Mean_gent-AKI +DMSO_=0.38 (N=26) vs. mean_gent-AKI +UPHD25_=0.15 (N=28). (D, E) Immunofluorescence co-stain of arginase-2 (red), macrophages (blue), and PT (green) in gent-AKI+DMSO (D) and gent-AKI +UPHD25 (E). Arg2+/mpeg1+ are marked with an asterisk and Arg-2-/mpeg1+ cells are marked with an arrow. (F) Quantification of M2 macrophage recruitment by counting Arg-2^+^/mpeg1^+^ cells adjacent to PT. Mean_gent-AKI +DMSO_= 0.61 (N=26) vs. mean _gent-AKI_+UPHD25=0.62 (N=27). Macrophage numbers were normalized by calculating ratio of M1 or M2 over total macrophages; i.e. (TNFα+/mpeg1+)/total mpeg1+. Data pooled from 3 biological replicates are shown expressed as mean+/-SEM. 2-tailed t-test: **p<0.01, n.s.= not significant. Scale bar=20 µM.

### Effect of PTBA on proximal tubular cell proliferation requires intact RA signaling

Among various pathways critical for macrophage response, the RA pathway has been implicated in macrophage recruitment during AKI (Chiba et al., 2015). Moreover, HDI treatments in mammalian AKI suppress inflammation and fibrosis, thereby improving long-term outcomes (Kinugasa et al., 2010; Marumo et al., 2009). Therefore, we determined whether PTBA required RA activity for efficacy in the zebrafish gent-AKI model. We treated gent-AKI *Tg(PT:EGFP)* fish with Ro41-5253 (abbreviated Ro41), an RAR antagonist that effectively blocks RA signaling in zebrafish larvae (Chiba et al., 2015). We analyzed proliferation by performing immunofluorescence and quantifying the number of PCNA+ RTECs. In DMSO treated larvae, Ro41-5253 did not significantly alter the number of PCNA+ RTECs compared to untreated larvae (Figure 6A,B,E). UPHD25 treatment increased the number of PCNA+ RTECs, while co-treatment with Ro41-5253 and UPHD25 significantly decreased PCNA+ RTECs compared to UPHD25 treatment alone (Figure 6C-E). Therefore, Ro41-5253 effectively blocks PTBA efficacy, suggesting that treatment requires intact RAR signaling to stimulate RTEC proliferation.

**Figure 6.**
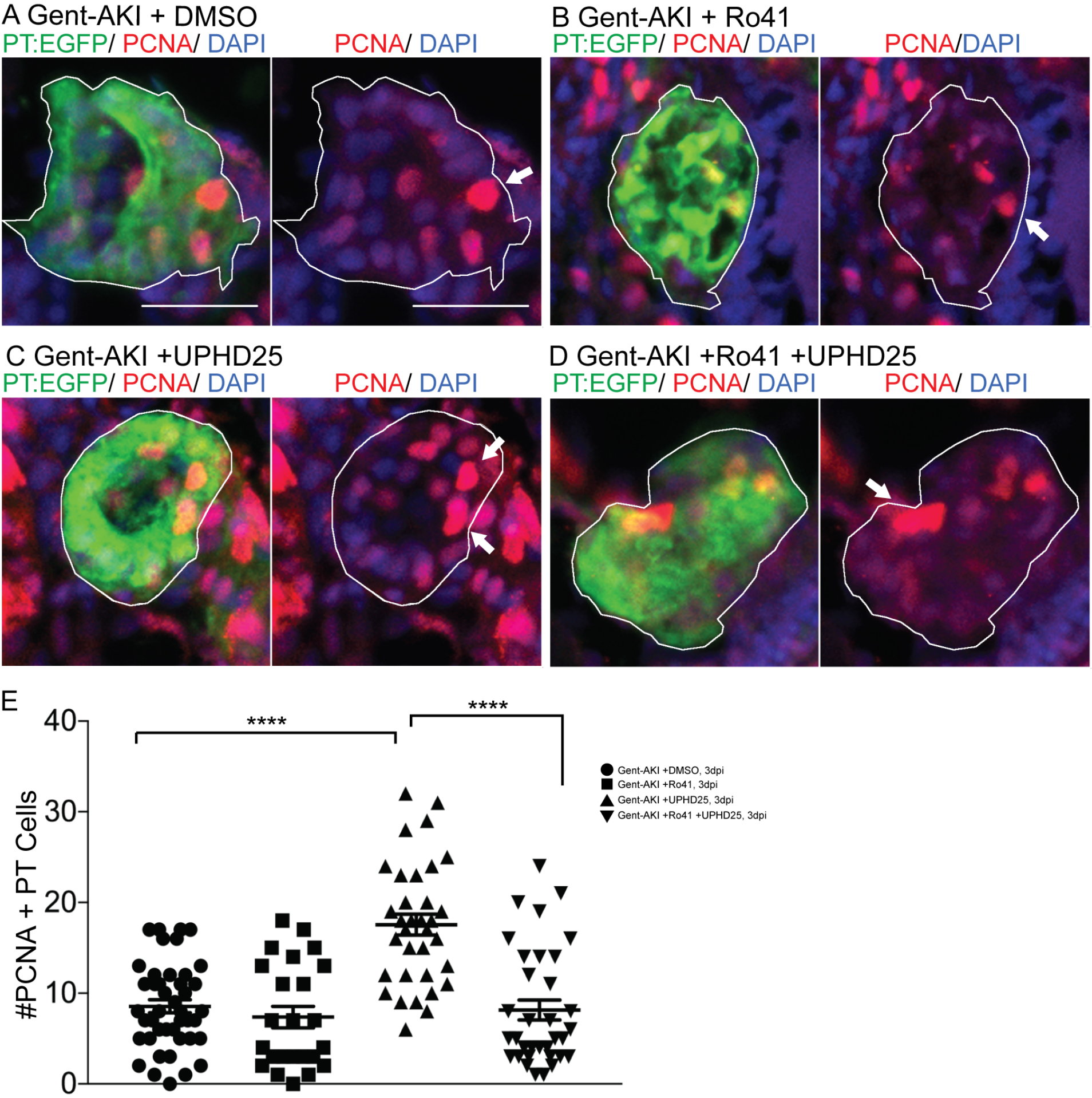
UPHD25 treatment efficacy requires intact RA signaling. *Tg(PT:EGFP)* transgenic zebrafish were used to analyze RTEC proliferation. (A-D) Immunofluorescence stain of actively undergoing S-phase marked with PCNA antibody (red) and PT (green) in gent-AKI +DMSO (A), gent-AKI +Ro41 (B), gent-AKI +UPHD25 (C), and gent-AKI +Ro41 +UPHD25 (D). PT is outlined in white and arrows mark PCNA+ RTECs. (E) Quantification of PCNA+ RTECs in each treatment group. Mean_UPHD25_=17.56 (N=34) vs. mean_Ro41 +UPHD25_=8.15 (N=34). Data pooled from 3 biological replicates are shown expressed as mean+/-SEM. One-way ANOVA: **p<0.01. Scale bar=20 µm.

To confirm the specificity of Ro41-5253 experiments, we utilized an inducible transgenic zebrafish line *Tg(hsp70l:EGFP-HS-dnRARα)* in which heat shock stimulates expression of a dominant-negative form of the human RA receptor alpha (DN-RARα)(Pogoda et al., 2018; Waxman et al., 2008; Waxman and Yelon, 2011). We previously showed that UPHD25 treatment increased active RTEC proliferation in uninjured larvae (Cianciolo Cosentino et al., 2013). Therefore, we evaluated whether reducing RA signaling in this heat-inducible transgenic model would block UPHD25-stimulated RTEC proliferation. Uninjured *Tg(hsp70l:EGFP-HS-dnRARα)* larvae were heat shocked for one hour at 5 dpf then treated with either DMSO or UPHD25 for 24 hrs. We quantified PCNA at 6 dpf and compared proliferation to wildtype larvae treated with DMSO or UPHD25 (Suppl. Figure 2A-E). Proliferation rates were comparable between heat-shocked (+HS) and non heat-shocked (-HS) larvae treated with DMSO (Suppl. Figure 2A,B); however, UPHD25 - HS treatment showed impaired RTEC proliferation compared to UPHD25 +HS (Suppl. Figure 2C,D). Therefore, in uninjured larvae, UPHD25 requires RA signaling to stimulate RTEC proliferation. Similarly, we evaluated proliferation in this transgenic model after AKI at 3 dpf. At 2 dpi, larvae were heat shocked for one hour and then treated with DMSO or UPHD25 for 24 hours (Suppl. Figure 2F-J). Overall, gent-AKI larvae showed higher proliferation than the no injury group, suggesting injury results in increased proliferation. Within the Gent-AKI group, proliferation rates were comparable between DMSO +HS and DMSO - HS (Suppl Figure 2F,G). However, gent-AKI +UPHD25 +HS showed impaired RTEC proliferation compared to UPHD25 - HS controls (Suppl. Figure 2H,I). Therefore, either pharmacologic or genetic inhibition of RAR signaling significantly reduces UPHD25 efficacy. Taken together, these data indicate that PTBA mechanism of action requires upstream RA signaling.

### PTBA enhances cardiac regeneration in adult zebrafish

Remote organ damage is one explanation for increased risk of morbidity and mortality in AKI (Depret et al., 2017). The heart, liver, lung, and brain are frequently damaged during AKI via several pathways, such as inflammatory cascades, apoptosis, and oxidative stress (Grams and Rabb, 2012; Nakazawa et al., 2017). Therefore, therapies that can target multiple organs will have greater therapeutic potential. To test PTBA’s efficacy as a multi-organ therapy, we evaluated treatment in an adult zebrafish model of cardiac injury (Poss et al., 2002), in which RA signaling has similarly been shown to be essential for regeneration (Kikuchi et al., 2011). Adult zebrafish recover from this procedure within 30-60 days and recovery relies on the regenerative capacity of proliferating cardiomyocytes, which peak around 7 days post amputation (dpa) (Jopling et al., 2010; Kikuchi et al., 2010; Poss et al., 2002). At 1 dpa, adult zebrafish were treated daily with 200 µM UPHD25 from 1-6 dpa (Missinato et al., 2015; Pugach et al., 2009). We evaluated cardiomyocyte proliferation at 7 dpa by staining for Mef2c, a marker of cardiomyocytes, and PCNA (Figure 7A-C). UPHD25 treatment did not affect cardiomyocyte proliferation in uninjured fish (Figure 7A,C), suggesting that PTBA does not affect non-injured, differentiated cells in adult tissue. After cardiac injury, UPHD25 treatment significantly increases the number of proliferating cardiomyocytes (mean_DMSO_=20.8 N=9 vs. mean_UPHD25_=34 N=8) (Figure 7B,C). To determine whether increased cardiomyocyte proliferation is associated with long-term beneficial effects, we performed acid fuchsin orange G (AFOG) staining after cardiac injury and quantified the fibrotic clot size at 20 dpa (Figure 7D). Compared to DMSO treated controls, UPHD25 treated fish showed a significant reduction in clot size, indicating acceleration of recovery (Figure 7D,E). Taken together, these data show that PTBA improves recovery after cardiac injury by increasing cardiomyocyte proliferation and decreasing fibrosis.

**Figure 7.**
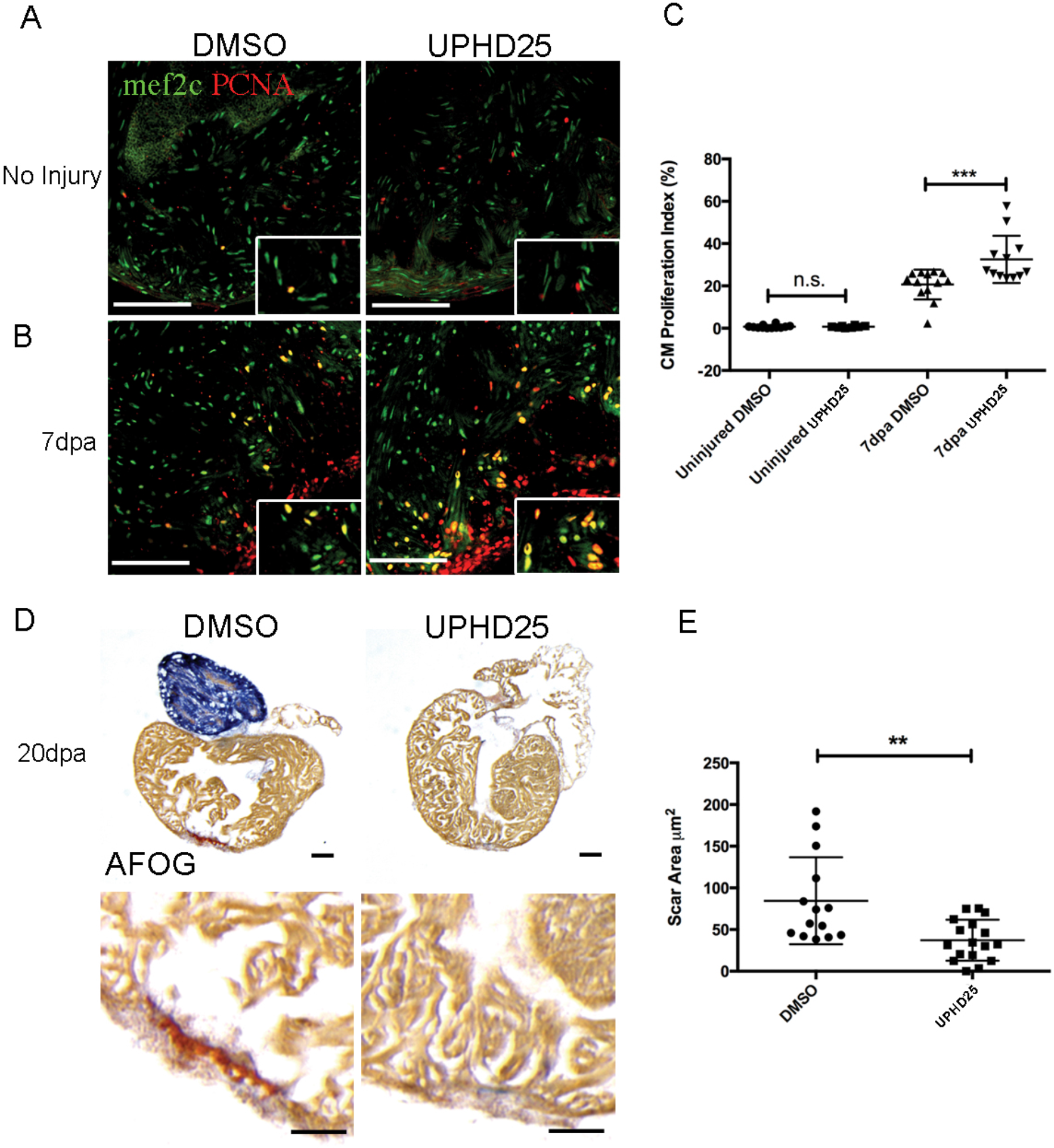
UPHD25 treatment ameliorates cardiac regeneration promoting cardiomyocytes proliferation. (A) Mef2c and PCNA immunostaining of uninjured hearts, injected for 6 days with DMSO (N=12), or 0.4 mg/Kg UPHD25 (N=8). (B) Heart at 7 dpa injected with DMSO (N=9) and UPHD25 (N=8). (C) Graph showing cardiomyocyte proliferation index in uninjured and 7 dpa hearts injected with UPHD25. (D) AFOG staining of hearts at 20 dpa injected with DMSO (N=14), and UPHD25 (N=17). Intact cardiac muscle is stained in yellow-orange, fibrin in red and collagen in blue. (E) Graph showing scar size at 20 dpa after injection of UPHD25. Mean_DMSO_=84.54 (N=14) vs. mean_UPHD25_=37.12 (N=17). Data pooled from 3 biological replicates are shown expressed as mean+/-SEM. 2-tailed t-test: ** P<0.01. n.s., not significant. Scale bars=100 µm.

## DISCUSSION

In this study, we showed that treatment with the PTBA prodrug UPHD25 enhances recovery after gent-AKI in zebrafish larvae by ameliorating tubular injury and stimulating RTEC dedifferentiation and proliferation. This work demonstrates PTBA is a useful chemical probe to dissect early post-repair processes in RTECs involving dedifferentiation and proliferation. PTBA treatment also increased cardiomyocyte proliferation following heart injury. Taken together, these studies indicate PTBA may possess broad therapeutic potential across multiple organ injury models.

There is evidence that cellular stress following injury results in cell-cycle arrest of RTECs (Yang et al., 2010). Prolonged cell-cycle arrest at either G1/S or G2/M checkpoints promotes maladaptive tubular responses associated with impaired repair and increased fibrosis after AKI (Ferenbach and Bonventre, 2015; Kellum and Chawla, 2015). We previously determined that PTBA drives an increase in the number of actively proliferating tubular epithelial cells and reduces the number of tubular epithelial cells in G2/M (Cianciolo Cosentino et al., 2013; Novitskaya et al., 2014). Since post-AKI fibrosis is thought to result from the accumulation of injured tubular epithelial cells in G2/M (Lin et al., 2010; Liu et al., 2013; Varmeh et al., 2011; Yang et al., 2010; Zahedi et al., 2006), these data suggest that PTBA reduces fibrosis by promoting effective tubular repair by more cells progressing through the G2/M checkpoint. Showing that PTBA induces an increase in Pax2a and Vimentin, known markers of RTEC dedifferentiation (Imgrund et al., 1999; Maeshima et al., 2002; Witzgall et al., 1994), further suggests the therapeutic effects of PTBA occur via an increase in the number of dedifferentiated RTECs with a high proliferative potential.

Mouse studies have shown that PTBA treatment enhances recovery from AKI by influencing the innate immune response. After treating with the nephrotoxin aristolochic acid, Novitskaya *et al*. observed an overall reduction in macrophage number that correlated with increased functional recovery and decreased fibrosis (Novitskaya et al., 2014). This is consistent with other reports of beneficial, anti-inflammatory properties of HDIs in kidney disease models (Venkatachalam et al., 2010; Wynn, 2010). In zebrafish gent-AKI, we saw a small, but significant difference in the number of early infiltrating macrophages after treatment. These findings suggest PTBA treatment may reduce the innate immune response either by promoting a healthier microenvironment or by altering the response from an inflammatory (M1) to repair (M2) environment. In agreement with promoting a favorable repair environment, treatment was found to reduce Kim-1 expression in injured nephrons and decrease the number of M1 macrophages, data that consistent with the PTBA driven response in mouse models of AKI (Cianciolo Cosentino et al., 2013; Novitskaya et al., 2014). No change in the number of M2 macrophages between treated and untreated larvae was observed, suggesting PTBA likely does not play a direct role in promoting macrophage polarization from pro-inflammatory to pro-repair. However, longitudinal studies may be needed to observe M2 conversion, as studies have shown monocytes can infiltrate into sites of acute injury and differentiate into M2 macrophages at later time points (Dey et al., 2014). Overall, these findings indicate that PTBA improves the environment early during injury by promoting dedifferentiation/proliferation of tubular cells and preventing excessive peritubular macrophage recruitment.

We demonstrated that the pro-proliferative effects of PTBA on RTECs requires RA signaling. Activation of the RA pathway in RTECs has been shown to be a critical early step during the regenerative response, and HDACs are known to repress RA signaling (Brilli et al., 2013; Chiba et al., 2015). Thus, it is possible that PTBA may lower the threshold concentration of RA required to initiate the activation of downstream targets or possibly prolong the window of active RA signaling (Menegola et al., 2006). Consistent with this idea, we showed that PTBA-enhanced proliferation only occurs in the context of injury and is dependent on intact RA signaling. This link between RA signaling and HDAC inhibition is best characterized in the cancer field, which often utilizes combinatorial RA-HDI chemotherapy as a potent treatment for malignancies that have shown resistance to RA therapy (Altucci and Gronemeyer, 2001; Berg et al., 1997; Bushue and Wan, 2010; Jopling et al., 2010; Pili et al., 2012). In cancer cells, HDIs alter gene expression and restore sensitivity to retinoid treatment (Jiang et al., 2008; Rettig et al., 2015; Touma et al., 2005; Wang et al., 2005). In our current study, we demonstrate that RA and HDI treatment can also work together during kidney regeneration. Interestingly, we found that PTBA treatment can enhance cardiac regeneration by promoting cardiomyoctye proliferation. Importantly, in uninjured hearts PTBA does not increase cell proliferation, but has an effect only after cardiac injury is performed. As with the kidney injury studies, we speculate that this is because RA production greatly increases in the zebrafish heart after injury to promote cardiomyocyte growth (Kikuchi et al., 2011).

Taken together, the current work provides a cellular mechanism of how PTBA accelerates renal recovery during AKI and cardiac regeneration. The goal of ongoing and future studies is to determine HDAC isoform selectivity and ultimately the molecular targets of PTBA. The most likely target appears to be class I HDACs. One attractive candidate is HDAC8, which has been linked to the repression of the developmentally important renal transcription factor gene *Lhx1* (Haberland et al., 2009; Saha et al., 2013). Furthermore, HDAC8 has been connected to RA signaling in cancer. Combined treatment of neuroblastoma cells with all-trans retinoic acid and an HDAC8 inhibitor resulted in enhanced tumor cell death (Rettig et al., 2015). Ultimately, studies to characterize the HDAC isoform selectivity and downstream molecular targets of PTBA will further our understanding of how this class of compound mitigates renal injury and advance efforts to develop a new drug therapy for AKI.

## MATERIALS AND METHODS

### Zebrafish husbandry

Studies were approved by the University of Pittsburgh IACUC. Zebrafish were maintained as described (Westerfield, 1993). In addition to AB wildtype, embryos were used from the following published transgenic lines: *Tg(PT:EGFP)nz4* (Cianciolo Cosentino et al., 2013), *Tg(cdh17:mCherry)pt307* (Chiba et al., 2015), *Tg(lyz:EGFP)nz117* (Kitaguchi et al., 2009), and *Tg(mpeg1:dendra2)uwm12* (Harvie et al., 2013), For *in vivo* imaging, larvae were kept in E3 medium containing 0.003% 1-pheny1-2-thiourea (PTU) after 24 hrs post fertilization.

### Generation of transgenic line Tg(hsp70l:EGFP-HS-dnRARa)

Tg*(hsp701:GFP-dn_Hsa.rarα)* transgenic fish were generated by gateway-based Tol2 transposon transgenesis(Kwan et al., 2007). To generate the transgenic fish, gateway cloning was used with the *hps70l* vector as the 5’ entry clone and GFP-dnRARa, a previously reported human dominant negative RARa, as the middle entry clone(Pogoda et al., 2018; Waxman and Yelon, 2011). PolyA was used as the 3’ entry clone. Following heat-shock induction, transgenic embryos expressed GFP and exhibited overt phenotypes, such as enlarged hearts and loss of forelimbs, which are consistent with loss of RA signaling(Waxman and Yelon, 2011) (data not shown). Constructs were injected into single cell embryos and screened for insertion. Progeny of P_0_ founder animals were used to establish the zebrafish transgenic line. In order to examine the effect of dnRAR during AKI, 2 dpi or 5dpf zebrafish larvae were heat shocked at 37°C for 1 hour. Then, the larvae were immediately treated with 1µM UPHD25 or 1% DMSO for 24 hours at 28°C. After treatment, the larvae were fixed and cryosectioned as described in the main methods. The cryosections were then immunostained with rabbit anti-PCNA antibody as described in the main methods.

### Gentamicin-induced AKI

Zebrafish larvae were injected with a single dose of gentamicin as previously described (Cianciolo Cosentino et al., 2010). Briefly, larvae were anesthetized in 160 mg/ml tricaine (Sigma-Aldrich)/E3 medium and injected with a total of 1 nl gentamicin (8-12ng) (Sigma-Aldrich) diluted in saline into the common cardinal vein. After injection, larvae were incubated in 50 µg/ml penicillin/streptomycin diluted in E3 medium.

### Cardiac injury

Adult AB* or Tu wildtype zebrafish age 6-18 months were anesthetized with 0.168 g/L tricaine for 3-5 minutes and ventricle apex amputation was performed as previously described (Missinato et al., 2018).

### Chemical treatments

All compounds were diluted in E3 medium containing 1% DMSO. UPHD25 was synthesized by Enamine and used at a working concentration of 1 µM. For RA inhibition studies, zebrafish larvae were treated with 1 µM Ro41-5253 (Enzo Life Sciences) in 1% DMSO diluted in E3 for 24 hrs from 3 to 4 dpf, and then compound was washed out with several changes of E3. In adults either 3 µL of 50% DMSO vehicle in PBS or 200 µM UPHD25 (corresponding to 0.4 mg/kg) was delivered by retro-orbital injection (Pugach et al., 2009). Injection was performed once per day, from 1-6 days post amputation (dpa), and hearts were extracted at 7 dpa to assess cardiomyocyte proliferation, and at 20 dpa to measure scar size.

### In situ hybridization

The *arginase-2* clone was synthesized and cloned into pEX-K248 with Sp6 promoter to drive the reverse transcription (Eurofins). The deoxygenin probe for *arginase-2* targeted 500 bp of 3’ end of coding region and 500 bp of 3’UTR. In situ hybridizations were performed as previously described(de Groh et al., 2010). Briefly, 6 dpf larvae were fixed in 4% paraformaldehyde (PFA)/PBS overnight at 4°C. Larvae were washed in PBS. Larvae were dehydrated in of MeOH and then moved to -20°C for 1 hr. Larvae were transferred to acetone for 10 min at -20°C. Larvae were rehydrated in PBS and treated with 100 ug/mL Proteinase-K in PBS/0.2%BSA/0.1%Tween-20 (PBTw) for 30 min at RT. Larvae were again fixed in 4% PFA for 20 min. Larvae were incubated with the *arg-2* probe overnight at 65°C. Larvae were incubated in 1:2000 anti-DIG Alkaline Phosphatase antibody (Roche) overnight at 4°C. The larvae were stained with BM purple (Roche) at room temperature. The reaction was stopped with 4% PFA. Larvae were then cryosectioned (see main methods) and imaged using a 20X objective on an Axiovert 40 CFL brightfield scope (Zeiss). Images were captured using Axiovision Rel v4.8 software (Zeiss).

### Histological analysis

Immunofluorescence microscopy was performed on cryosections as described previously (Drummond and Davidson, 2010). Larvae were fixed in 4% paraformaldehyde and treated with a 10-30% sucrose/PBS gradient before embedding in tissue freezing medium (Ted Pella). Sections were generated at a thickness of 12-14 µm. Slides were blocked with 10% goat serum in PBST (0.1% Tween-20), followed by primary and secondary antibody incubations (Supplemental Table 1). Incubation with DAPI (Vector Laboratories) was used to counterstain nuclei. Sections were examined by confocal microscopy (Zeiss LSM 700). The slides were washed with PBS then mounted with Aqua Polymount (Polysciences, 18606-20).

To quantify Pax2a-positive cells in *Tg(PT:EGFP)* fish, we imported serial images into ImageJ 1.46r software (NIH). The ROI tool was used to outline the kidney in the green channel, and “Analyze Particle” function was used to quantify the number of Pax2a-positive cells in the red channel. Background and threshold values were constant between groups for each experiment. Particle size range was 80 to infinity. Per nephron, three images were analyzed. Similar ImageJ analysis was performed in *Tg(PT:EGFP); Tg(mpeg1:dendra2)* fish to quantify macrophage cell size. For these images, there was no background removal, mpeg1:dendra2 channel threshold was 53 to 255, and particle size range was 60 to infinity.

Heart cryosections were stained with AFOG as previously described (Missinato et al., 2018). Images were captured with Leica MZ 16 microscope and Q Imaging Retige 1300 camera. Clot area was measured using ImageJ (NIH). Four heart sections showing the largest clot were measured and the scar area values were measured as average of the sum of the clot area (Supplemental Table 1). Slides were mounted in Vectashield with DAPI (Vector Laboratories). For each experiment, at least four sections were analyzed per heart. Cardiomyocyte proliferation index was calculated as percentage of number of Mef2c+ and PCNA+ cells divided by the number of total Mef2c+ cells.

### Live confocal zebrafish imaging

24 hrs after gentamicin injection, *Tg(chd17:mCherry);Tg(lyz:EGFP)* larvae were anesthetized in tricaine, embedded in a thin layer of 0.5% low-melt Sea Plaque agarose (Cambrex), and covered with E3 medium plus PTU to prevent pigment development. Image stacks were acquired using a Leica TCS SP5 multiphoton microscope (Leica Microsystems, Wetzlar, Germany) with an HCX IRAPO L 20X/0.95 water immersion objective, non-descanned detectors and a custom built motorized stage (Scientifica, East Sussex, UK). Sequential stack scanning was performed bidirectionally with a resonant scanner (16000 Hz, phase set to 1.69) with 32x line averaging and a zoom of 1.7x. EGFP and mCherry were excited with a Mai Tai DeepSee Ti:Sapphire laser (Newport/Spectra Physics, Santa Clara, CA) at 900 and 561 nm, respectively. Using the “Mark and Find” function, (x,y) coordinates and z-series parameters (step size 1.48 µm) were defined for individual larvae. Images were captured every 27 minutes for 17 hrs. Maximal projections were compiled in series to generate time-lapse movies using LAS AF Version: 3.0.0 build 8134 and Metamorph software.

48 hrs after gentamicin injection, *Tg(chd17:mCherry);Tg(mpeg1:dendra2)* larvae were processed for imaging using the same protocol documented above. Image stacks were acquired using Nikon Eclipse Ti confocal microscope (Nikon Instruments, Melville, NY, USA) with a 20X dry objective, and a motorized stage. Stacks were captured with 40 optical sections, with 5 µm step size. Dendra2 and mCherry were excited with 488nm and 560nm lasers, respectively. Dendra2 is photoconvertible with 488nm and UV but maintains its original green fluorescence emission (509nm) when imaged under laser power between 2-7%. Experiments utilizing *Tg(mpeg1:dendra2)* maintained low laser power to inhibit photoconversion from green to red. Using the “Mark and Find” function, (x,y) coordinates and z-series parameters (step size 5 µm) were defined for individual larvae. Images were captured every 90 minutes for 24 hrs. Maximal projections were compiled in series to generate time-lapse movies using Imaris image analysis software (Bitplane, Zurich, Switzerland).

### Statistical analysis

Data were analyzed using student t-test, one-way ANOVA, and two-way ANOVA as indicated, and data are reported as mean ±SEM. P values were considered significant when <0.05. For studies in zebrafish larvae, “N” reflects the number of nephrons included in the analysis per group. When visible, both nephrons per fish were included. For adult zebrafish studies, “N” reflects the number of hearts included per group.

## ACKNOWLEDGMENTS

The Hukriede laboratory is supported by the National Institutes of Health (NIH) National Institute of Diabetes, Digestive and Kidney Diseases (NIDDK) grants 2R01DK069403, 1R01DK112652, 1P30DK079307, the Eunice Kennedy Shriver National Institute of Child Health and Human Development (NICHD) grant 2R01HD053287, and the Department of Defense W81XWH-17-1-0610. LBS was supported by F30DK101143. EE was supported by T32DK061296. The Davidson laboratory is support by the Health Research Council of New Zealand (grants 15/057 & 17/425). The Tsang laboratory is supported by grants from the NIH (R01HD053287) and AHA (14GRNT20480183).

## AUTHOR CONTRIBUTIONS

L.B.S., H.I.H., E.B.E., M.A.M., J.S.W., S.C.W., A.J.D., M.T., and N.A.H. designed experiments;L.B.S., H.I.H., E.B.E., M.A.M., M.D.M., E.R.R., J.S.W., and S.C.W. performed experiments;L.B.S., H.I.H., E.B.E., M.A.M., M.D.M., E.R.R., B.L.R., J.S.W. S.C.W., A.J.D., M.T., and N.A.H.analyzed and interpreted data. L.B.S., H.I.H., M.A.M., A.J.D., M.T., and N.A.H. wrote the paper.

## DISCLOSURES

All authors declared no competing interests.

